# Pseudogenisation of NK3 Homeobox 2 (*Nkx3.2*) in Monotremes Provides Insight into Unique Gastric Anatomy and Physiology

**DOI:** 10.1101/2024.03.13.584895

**Authors:** Jackson Dann, Zhipeng Qu, Linda Shearwin-Whyatt, Rachel van der Ploeg, Frank Grützner

**Affiliations:** School of Biological Sciences, The University of Adelaide, SA 5005, Australia

## Abstract

Development of the vertebrate antral stomach and pyloric sphincter (antropyloric region) – involved in enzymatic breakdown and thoroughfare of food - is underpinned by a highly conserved developmental pathway involving the hedgehog, bone morphogenetic protein (BMP) and Wingless/Int-1 (Wnt) protein families. Monotremes are a unique lineage where acid-based digestion has been lost, and this correlates with a lack of genes for gastric acid and enzymes in the genomes of the platypus (*Ornithorhynchus anatinus*) and short-beaked echidna (*Tachyglossus aculeatus*). Furthermore, these species feature unique gastric phenotypes, both with truncated and aglandular antral stomachs and the platypus with no pylorus. Here, we explore the genetic underpinning of monotreme gastric phenotypes, investigating genes important in antropyloric development using the newest monotreme genome sequences (mOrnAna1.pri.v4 and mTacAcu1) together with RNA-seq data. We found that the pathway is generally conserved but, NK3 homeobox 2 (*Nkx3.2*) was pseudogenised in both platypus and echidna. We speculate that pyloric-like restriction in the echidna may correlate with independent evolution of *Grem1* and *Bmp4* sequences, and that the convergent loss of gastric acid and stomach size genotypes and phenotypes in teleost and monotreme lineages may be a result of eco-evolutionary dynamics. These findings reflect the effects of gene loss on phenotypic evolution and further elucidate the genetic control of monotreme stomach anatomy and physiology.

## Introduction

Monotremes (order Monotremata) are a unique mammalian phylum in their gastric anatomy and physiology which diverged from therian mammals 187 million years ago. The platypus (*Ornithorhynchus anatinus*) and the short-beaked echidna (*Tachyglossus aculeatus*) are two of the few vertebrate species to have lost acidic gastric juices and glandular antral epithelium^1^. Changes in diet and ecological specialisation since their divergence 55 million years ago have led to morphological evolution of the gastrointestinal tract between these two monotreme species. The platypus stomach is small, amorphic and glandless and it lacks a pyloric sphincter, making it notably hard to distinguish from the oesophagus and intestines. The echidna stomach, whilst also glandless and non-acidic, is bulbous containing a pyloric-like restriction through to the duodenum to regulate the flow of food and gastric juices^2,3,4^.

The antral stomach and pyloric sphincter (antropyloric region) are shared features throughout vertebrate evolution which can be dated back to early diverging jawed-vertebrate lineages around 450 million years ago. Monotremes are one of the few lineages that have lost these structures, making them interesting species to investigate the molecular signatures associated with this change in digestive anatomy. The glandular antral stomach is responsible for secretion of hydrochloric acid to maintain the acidic luminal environment whilst the pyloric sphincter regulates the passage of food into the duodenum and prevents acidic juices from damaging duodenal epithelium. The development of this structure is controlled by a highly conserved developmental pathway found in zebrafish, chicken, frog and mouse^5,6^.

A key gene of this pathway, the NK3 homeobox 2 (*Nkx3.2*), has epithelial and mesenchymal roles in antropyloric development as well as other roles in axial and peripheral skeletal development^7,8^. During stomach development, *Nkx3.2* expression in undifferentiated gastric mesenchyme demarcates the future pylorus from the antral stomach and duodenum by repressing *Bmp4* expression and potentiating *Nkx2.5*, *Sox9*, *Six2* and *Grem1* expression. The *Barx1*-*Nkx3.2* signaling cascade at this timepoint also represses Wnt signaling (especially *Wnt5a*) to promote antral stomach identity^7,8,9^. Development of gastric epithelial cell identities such as gastrin-releasing G cells in the antral stomach and pylorus are regulated by interaction of sonic hedgehog protein (encoded by *Shh*) and *Nkx3.*2. Other proteins such as Indian hedgehog protein (*Ihh*), nuclear receptor subfamily 2 group F member 2 (*Nr2f2*) and pancreatic-and-duodenal homeobox 1 (Pdx1) potentiate epithelial cell type development using many additional pathways (Figure 1)^10,11,12^.

**Figure 1:**
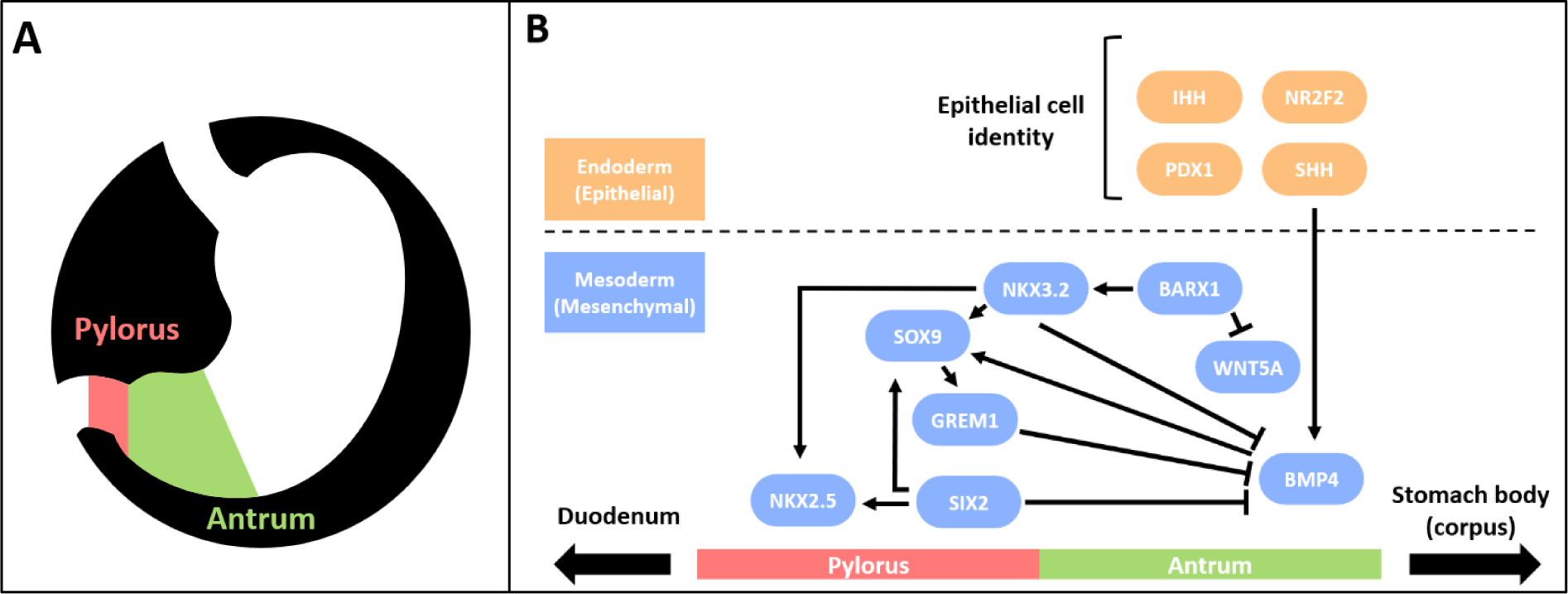
Diagrammatic representation of the stomach anatomy (A) and the conserved vertebrate antropyloric developmental pathway (B). Constituents of the pathway are divided according to their roles in epithelial and mesenchymal development (orange and blue in B) and their positioning along the horizontal axis depicts whether they are implicated in pyloric and/or antral development. Figure adapted from Self et al. (2009) and Udager et al. (2010).

Here, we present the expression of antropyloric developmental genes in monotremes and conservation in a range of vertebrates, and identify *Nkx3.2* to have been pseudogenised in the monotreme ancestor. We then speculate further on what changes may be associated with morphological differences between the platypus and echidna and outline overlapping eco-evolutionary dynamics between monotremes and other vertebrate lineages which have lost gastric acid digestion.

## Materials and Methods

### Sequence retrieval, alignments and phylogenies

Protein products of *Nkx3.2*, *Barx1, Shh*, *Ihh, Pdx1, Nr2f2, Nkx2.5, Six2, Bmp4, Sox9, Grem1* and *Wnt5a* were acquired through NCBI GenBank (https://www.ncbi.nlm.nih.gov/genbank/). Sequences were obtained from eutherian mammals (humans / *Homo sapiens*, house mouse / *Mus musculus*, brown rat / *Rattus norvegicus*, horse / *Equus caballus*, cows / *Bos taurus* and pig / *Sus scrofa*); marsupials (common wombat / *Vombatus ursinus* and gracile agile opossum / *Gracilinanus agilis*); reptiles (chicken / *Gallus gallus*, fence lizard / *Sceloporus undulatus*), bony fishes (zebrafish / *Danio rerio*) and cartilaginous fishes (salmon / *Salmo salar*). Sequences for monotremes (platypus / *Ornithorhynchus anatinus* and short-beaked echidna / *Tachyglossus aculeatus*) were acquired from the newly-released genome sequences: mOrnAna1.pri.v4 and mTacAcu1.pri respectively^13^.

Multiple sequence alignments of derived protein sequences were performed using the Clustal Omega plugin in Geneious Prime 11.0.14+1^14,15^. Protein phylogenies were constructed using the online IQ-TREE web server with ModelFinder for substitution model and UFBOOT for 1000 bootstrap replicates. Amino acid substitution models were chosen based on Bayesian information criterion maximum log likelihood estimates^16,17,18^. Substitutions, insertions and deletions occurring in conserved domains that occurred exclusively in one or both of the monotreme species were then noted and compared with associated literature or the conserved domain database (CDD)^19^.

### Expression analysis

RNA-Seq fastq files for platypus and echidna tissues were mapped to the platypus genome mOrnAna1.pri.v4 and mTacAcu1.pri using HISAT2^13,20,21,22^. Gene expression for *Nkx3.2* and other pathway genes were analysed using FeatureCounts 2.0.0 and RSEM 1.3.3 with default settings respectively. For *Nkx3.2* expression analysis exon coordinates were manually annotated in GTF format and random intergenic regions were chosen as a negative control for transcriptional noise^23,24,25^.

### gDNA/RNA preparation and cDNA synthesis

Snap frozen monotreme liver, stomach and spleen samples stored at -80°C were lysed using liquid nitrogen in a mortar and pestle. Monotreme liver gDNA was prepared using the phenol-chloroform method^26^. RNA samples from monotreme stomach, liver and spleen were prepared using the Qiagen RNeasy plus micro kit (Qiagen, Hilden, Germany) as per instructions. Total RNA samples from monotreme tissues were reverse transcribed using the iScript cDNA synthesis kit (Bio-Rad, Hercules, CA, USA). RNA was stored in nuclease free water at -80°C.

### PCR

Beta-actin (*Actb*) and NK3 homeobox 2 (*Nkx3.2*) primers were designed using the NCBI primer-blast (https://www.ncbi.nlm.nih.gov/tools/primer-blast/) and synthesised by integrated DNA technologies (Coralville, IA, USA), (Table 1).

**Table 1:**
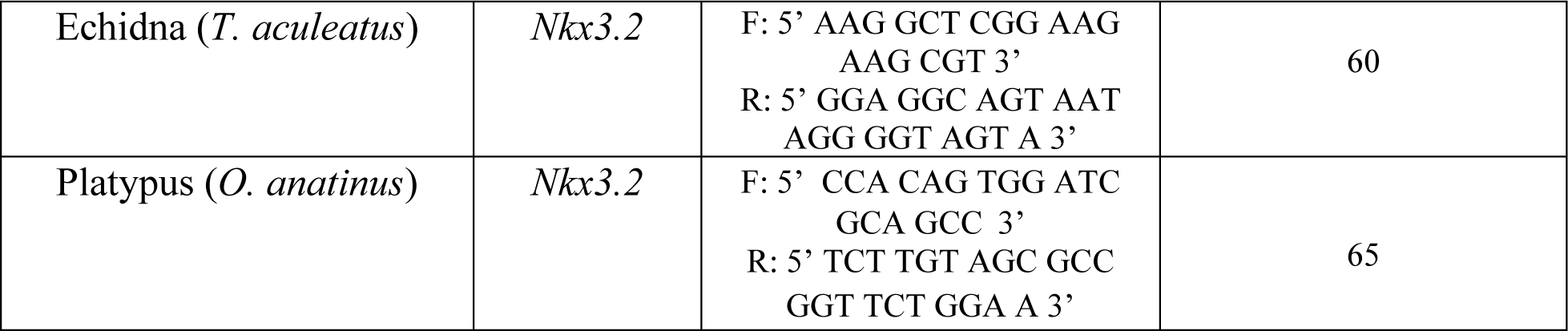
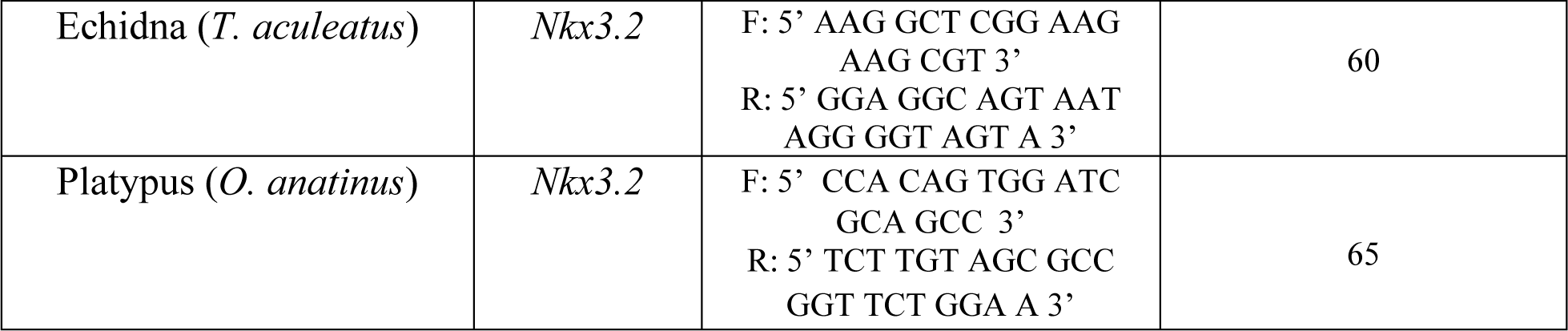
Species, genes and primer sequences used for gDNA PCRs and RT-PCRs.

Genomic DNA and cDNA PCRs were carried out using OneTaq polymerase (New England Biolabs, Ipswich, MA, USA) as per instructions. Each 25 μL reaction contained 5 μL GC reaction buffer, 0.5 μL 10 mM dNTPs, 1 μL forward / reverse primer 10 μM mix, 0.25 μL polymerase and 18.25 μL nuclease-free water. Cycling occurred in the C1000 touch thermal cycler (Bio-Rad, CA, USA). Conditions involved an initial denaturation at 94 degrees for 3 minutes, then 35 cycles of 30 seconds denaturation at 94 degrees, 30 seconds annealing at appropriate temperatures (Table 1) and extension at 68 degrees for 1 min/kb. All PCR products were confirmed by Sanger sequencing (Australian Genome Research Facility, Melbourne, VIC, AUS).

## Results

### Sequence changes and expression of antropyloric developmental genes

To determine the conservation of epithelial and mesenchymal antropyloric developmental genes, monotreme sequences were compared to those of a range of vertebrate species. Gene expression in monotremes was investigated using RNA-seq datasets and PCR products compared either with non-transcribed region controls or benchmark TPM standards (Table S1).

Of the epithelial genes, conserved domains of *Nr2f2*, *Pdx1*, *Shh* and *Ihh* monotreme amino acid sequences contained no changes likely to affect function and sequence identities were consistent with vertebrate species. All genes were expressed in at least one tissue beyond a significant threshold (> 0.5 TPM) except *Pdx1* in platypus, where no expression was detected (Table S1). In echidna *Pdx1* missense mutations entirely change the conserved D/EPEQD motif-responsible for PCIF-1 interaction -to GAPPGL ^27,28,29^. The Sonic Hedgehog protein sequences (SHH) contain an insertion of around 45 amino acids in the Hint domain and deletions of 24 and 34 amino acids at the C-terminal domain in the platypus and echidna respectively. The monotreme Indian Hedgehog sequences (IHH) contain insertions of 70 and 25 amino acids in the N-terminal signaling domain of echidna and platypus respectively (Table S1)^30^.

No major changes were seen in conserved domains and sequence identities outside of vertebrate ranges for the following mesenchymal genes: *Sox9*, *Six2*, *Wnt5a* and *Barx1*^31,32,33,34^. Gene expression was monitored in tissues using available RNAseq datasets (Table S1). Monotreme *Nkx2.5* sequences contained a variety of insertions and deletions in between conserved domains and missense mutations in the TD and NK-SD domains (Table S1)^35^. Monotreme *Bmp4* sequences contained several short (9 to 15 amino acids) insertions N-terminal to- and within the signaling domain respectively and 7 to 14 amino acid insertions N-terminal to the second splice site^36^. The echidna *Grem1*-derived protein sequence contained an N-terminal insertion of 16 amino acids, and both sequences shared several missense mutations in the *Bmp4* inhibitory domain not conserved elsewhere^37^.

### Vestigial Nkx3.2 sequences remain in the conserved syntenic region in monotremes

As a key gene implicated in epithelial and mesenchymal development of the antropyloric region, we sought to examine its conservation and expression in monotreme species^5^. Using conserved vertebrate domains from Lettice, Hecksher-Sørensen and Hill^38^, some sequence remnants were identified in monotreme genomes. To ensure the sequences identified were asociate with *Nkx3.2*, synteny analysis was performed. The three genes flanking either side of *Nkx3.2* are entirely conserved with the exception of the fence lizard (*Sceloporus undulatus*) which is missing CLNK at the terminus of the region (Figure 2). Reconstructed monotreme *Nkx3.2* sequences occurred in the conserved syntenic region between *Rab28* and *Bod1l1*. Given a lack of annotations, intron-exon boundaries were then estimated using conserved vertebrate motifs from Lettice, Hecksher-Sørensen and Hill^38^, and average conserved exon lengths from other mammalian taxa (Figure S1).

**Figure 2:**
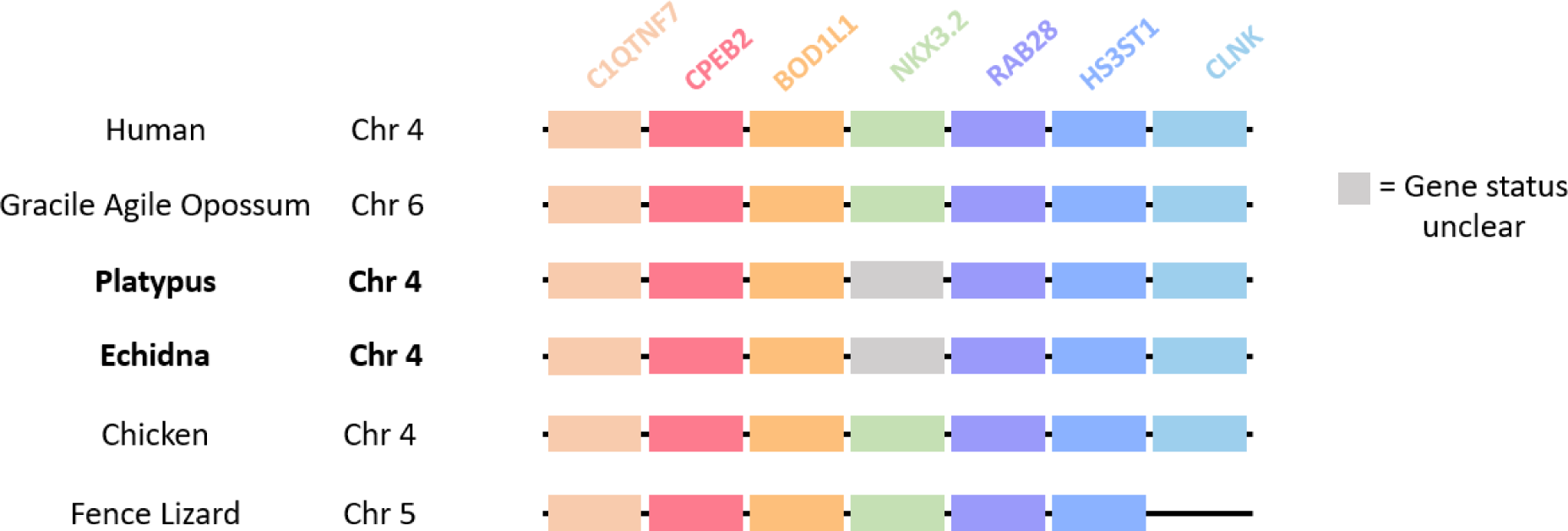
Syntenic analysis of the genomic neighbourhood of *Nkx3.2* in human (*Homo sapiens*, chr.4: 10,480,039-13,631,835), gracile agile opossum (*Gracilinanus agilis*, chr6: 203,879,610-209,238022), platypuse (*Ornithorhynchus anatinus*, chr.4: 120,552,553-125,119,742), echidna (*Tachyglossus aculeatus*, chr4: 12,899,770-18,239904), chicken (*Gallus gallus*, chr4: 76,333,940-77,775,864) and fence lizard (*Sceloporus undulatus*, chr5: 95,631,178-96,899,783).

Monotreme *Nkx3.2* sequences were then translated and aligned to determine significant sequence changes in conserved domains (Figure 3). The identities of reconstructed sequences ranged between 37-55% when compared with those of vertebrates, which was lower than other vertebrate sequences with each other; ranging from 50-95% sequence identity. The lower-than-expected sequence identities in monotremes may be attributed to an accumulation of various insertions/deletions and mutations in between the C-terminal box, homeobox and N-terminal box. These conserved domains in NKX superfamily proteins were mostly conserved between monotreme species. After reconstructing the open reading frame using conserved vertebrate intron-exon boundaries the echidna sequence contained a premature stop codon (Figure 3).

**Figure 3:**
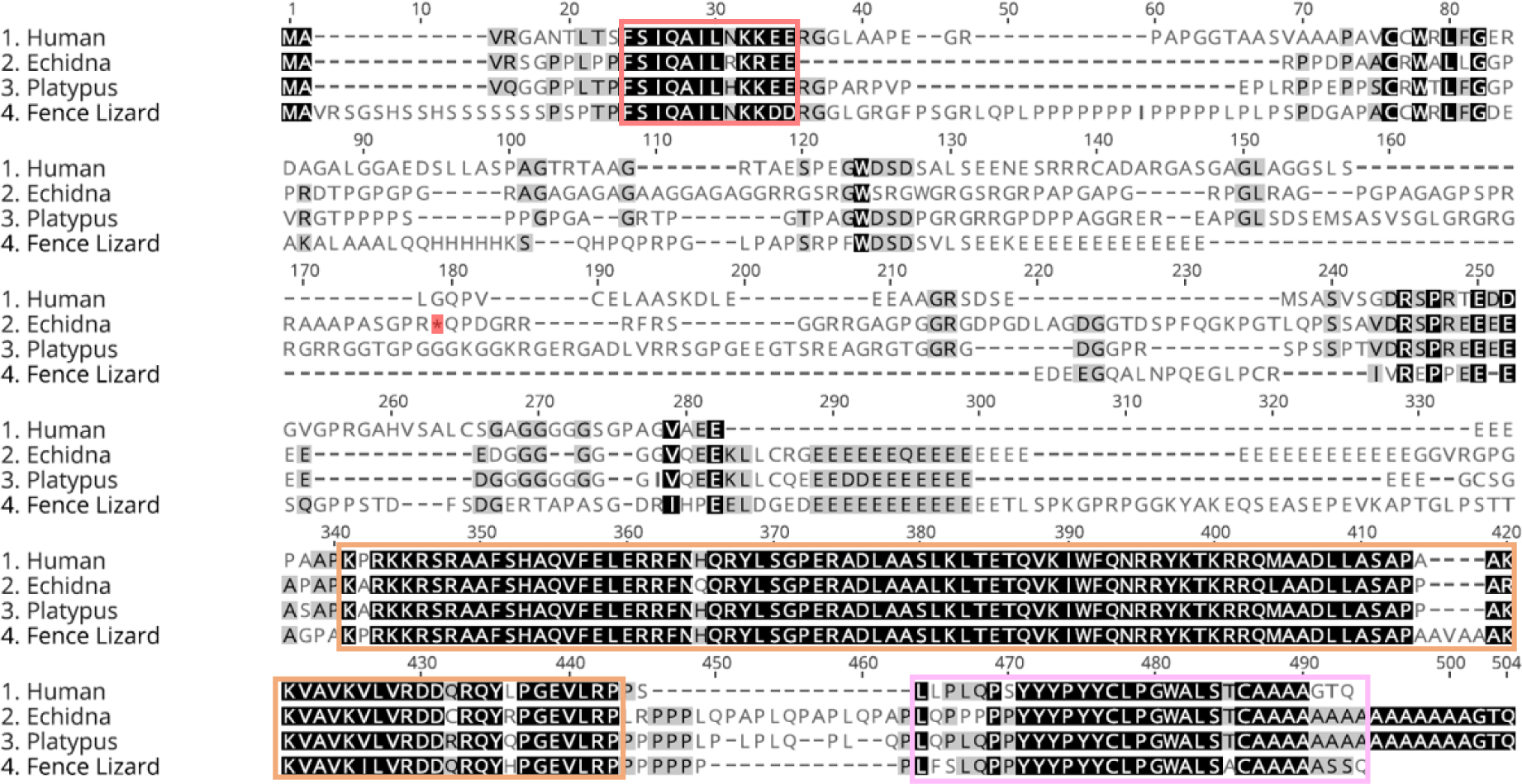
Geneious Clustal Omega Alignment of NKX3-2 amino acid sequences from humans (*Homo sapiens*), short-beaked echidnas (*Tachyglossus aculeatus*), platypuses (*Ornithorhynchus anatinus*) and fence lizards (*Sceloporus undulatus*) outlining the accumulation of mutations, insertions and deletions in the monotreme sequences. Residues in the red box form the C-terminal box, orange the homeobox and pink the N-terminal box all conserved domains of the NKX protein superfamily. A premature stop in the echidna amino acid sequence is highlighted in red and darker shading of residues indicates higher sequence similarity.

### Monotreme Nkx3.2 shows no significant expression in adult and juvenile monotreme tissues

Major sequence changes in *Nkx3.2* suggest that this gene has mutated extensively so we investigated its transcriptional activity. The *Nkx3.2* exon coordinates did not align with any liver RNA-seq data in NCBI, so expression was probed further using RT-PCR. Few reads mapped to the predicted exons of *Nkx3.2* from a variety of platypus and echidna tissues (Table 2) and most reads mapped could be corroborated with intergenic controls therefore transient non-significant expression (Table 3).

**Table 2:** Number of reads mapped to exon coordinates for *Nkx3.2* in various tissues from the two monotreme species and the resulting RPKM values.

|  | Tissue | Mapped read total for gene | Mapped read total for tissue | RPKM |
| --- | --- | --- | --- | --- |
| Echidna | Adult Ovary | 0 | 54548375 | 0 |
|  | Adult Testes | 0 | 61299876 | 0 |
|  | Adult Frontal Cortex | 0 | 58422726 | 0 |
|  | Puggle Frontal Cortex | 0 | 47214854 | 0 |
|  | Puggle Ovary | 2 | 47923676 | 0.007713 |
|  | Puggle Testes | 0 | 54413890 | 0 |
| Platypus | Brain | 0 | 37686965 | 0 |
|  | Fibroblasts | 0 | 39841608 | 0 |
|  | Kidney | 0 | 18380826 | 0 |
|  | Liver | 0 | 38897053 | 0 |
|  | Ovary | 1 | 25060266 | 0.012222 |
|  | Testes | 0 | 66530964 | 0 |

**Table 3:**
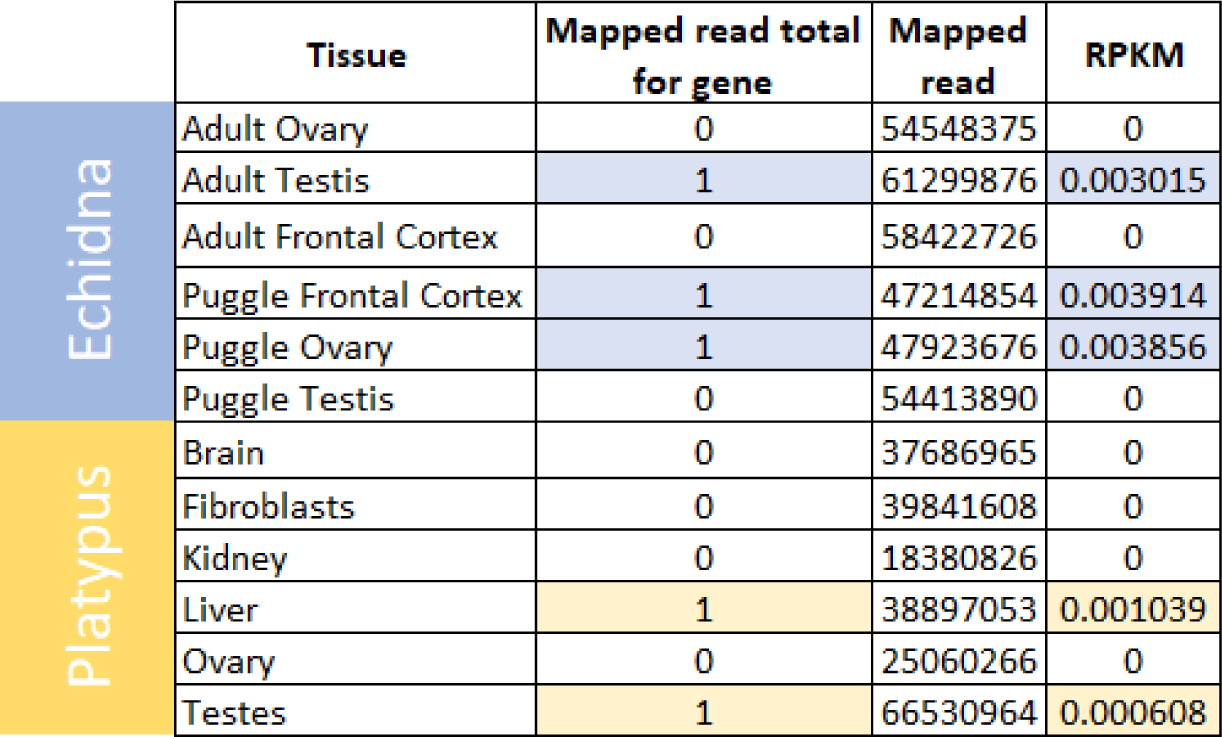
Number of reads mapped to randomly selected non-transcribed regions in various tissues from the two monotreme species and the resulting RPKM values.

To verify the absence of expression, RT-PCR was carried out on platypus and echidna stomach, spleen and liver tissues. Β-actin genomic DNA and expression was used as controls from each tissue. No expression was observed (Figure 4). We conclude that monotremes have lost a functional *Nkx3.2* gene.

**Figure 4:**
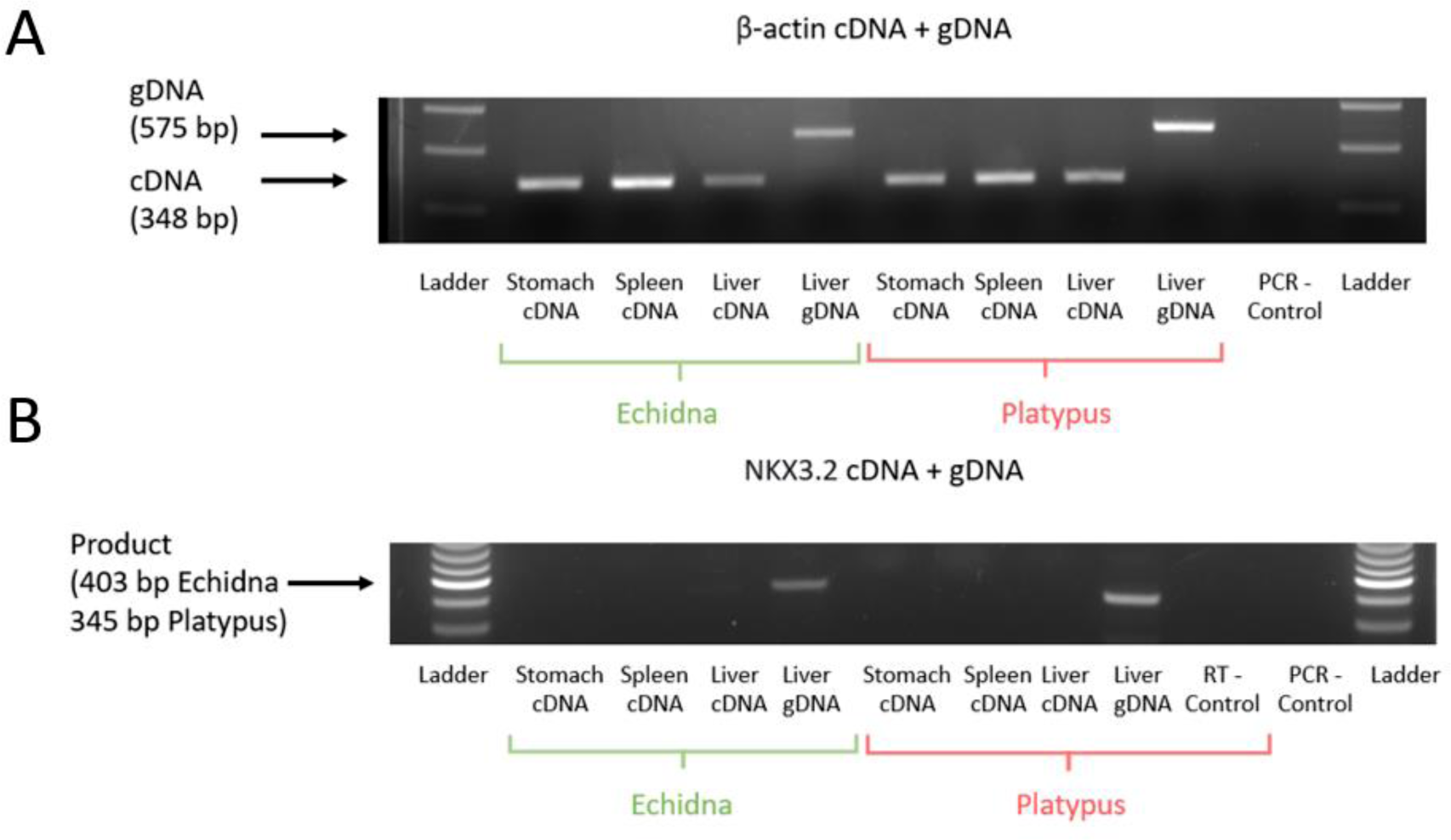
Genomic and RT-PCRs of platypus and echidna cDNA and gDNA from stomach, liver and spleen using primers for β-actin (A) and NKX3.2 (B). Due to high-GC content and large introns, intra-exonic primers were used for *Nkx3.2* PCRs where absence vs presence would be indicative of expression.

## Discussion

Since their divergence from therian mammals 187 million years ago, gastrointestinal morphology has changed significantly in echidna and platypus. There has been a loss of acidic gastric juices, antral glandular epithelium, a reduction in stomach size in the platypus and a loss of the pylorus in the platypus despite the echidna displaying a pyloric-like restriction^1,2,5^. We investigated the highly conserved vertebrate antropyloric developmental pathway to determine the genetic basis for these contrasting phenotypes.

### Sequence changes in antropyloric developmental genes provide clues for gastric phenotype discrepancy

Overall we found conservation of the antropyloric developmental genes *Nr2f2*, *Sox9*, *Six2*, *Wnt5a* and *Barx1*. Insertions, deletions and missense mutations in *Shh* and *Nkx2.5* were mostly conserved between monotreme species and therefore unlikely to be associated with the different gastric phenotypes. Of the epithelial genes, echidna *Pdx1* was missing the majority of the canonical PCIF-1 domain sequence. The PCIF-1 domain allows the PDX-1-PCIF-1 interface and transcriptional inhibition of PDX-1 which is necessary for pancreatic islet and duodenal development^31^. Expression of *Pdx1* was close to the significant threshold (0.5 FPKM) in echidna ovaries and testis but otherwise not present in transcriptome data, and synteny of the *Pdx1* region was conserved. The echidna Indian hedgehog (*Ihh*) sequence contained a large insertion of 70 amino acids in the n-terminal signalling domain, compared to a 25 amino acids insertion in the platypus. This domain is necessary for transduction of the hedgehog signalling pathway^30^. Of the mesenchymal genes, the echidna *Grem1* contained a unique N-terminal insertion in the signal peptide and sequence change in the BMP inhibitory domain shared with the platypus. The inhibitory domain prevents BMP-receptor binding and the downstream signalling pathway, a necessary step in regulation of many fundamental developmental processes^36^. Echidna *Bmp4* also contained a unique missense mutation at the furin S2 cleavage site, which probably precludes activity and normal signaling of the mature peptide^37^.

Surprisingly, our results show that *Nkx3.2* is not expressed in either species and the genes have accumulated neutral mutations and a stop codon which is likely to reflect pseudogenisation. This suggests a pseudogenisation event in the most recent ancestor of monotremes that may correlate with previously noted deletions of gastric enzyme genes and those associated with hydrochloric acid production.

### The Nkx3.2 pseudogenisation likely occurred in the monotreme ancestor and is probably associated with the loss of antral and pyloric segment identities

Transcriptome and RT-PCR results from this study showed an absence of *Nkx3.2* expression in any monotreme tissues. This together with an accumulation of missense mutations not found elsewhere and a predicted premature stop codon in echidna, provide strong evidence for a shared pseudogenisation that occurred in the monotreme ancestor before their divergence 55 million years ago. Pseudogenisation of *Nkx3.2* in monotremes and their corresponding gastric phenotypes mirrors the phenotype of *Nkx3.2* knockout mice which show no pyloric restriction, truncated antral sections and markers for antral glands were severely reduced and located near the duodenum or lost^39,40^.

In the absence of *Nkx3.2*, establishment of the echidna pyloric-like restriction would likely require compensatory inhibition of *Bmp4* as is seen in mouse and chicken developmental models^4^. Knockout *Six2* and *Nkx3.2* models show the additive and essential roles of both these genes in boundary formation, with *Grem1* misexpression studies showing additive inhibition of *Bmp4* expression but no effect on the pyloric phenotype^40^. Of the sequence changes listed above, the most likely candidates which may cause the gastric phenotypic difference between the platypus and echidna are *Grem1* and *Bmp4* as they both affect the *Bmp4* expression domain and therefore establishment of the boundary between the pylorus and antral stomach. The insertion in the signal peptide domain of echidna *Grem1* would only affect protein translocation and secretion, which may have no effect on *Bmp4* inhibitory capacity. The missense mutations in the BMP inhibitory domain of *Grem1* may affect protein-protein interactions but are shared between monotreme species. In echidna *Bmp4*, the missense mutation in the furin S2 cleavage site may affect activity and signaling but it is unclear whether this would also affect other aspects of development given the broad function of the protein.

### Correlations between Nkx3.2 pseudogenisation and gastric gene loss in the monotreme ancestor

The antral glandular portion of the mammalian stomach contains the vast majority of G cells, the endocrine cells responsible for gastrin release, hydrochloric acid secretion, mucosal growth and stomach motility^41,42,43^. Transcriptional activity of gastrin-associated genes in mice and humans reflect the importance of this region for acid secretion, with over 90% occurring in these units^44,45^. In monotremes, however, these genes have been pseudogenised, along with those encoding pepsinogen A and C, the sodium/potassium ATPase pump proteins and cathepsin E^3,4^. These genes represent the necessary constituents for maintenance of the acidic luminal environment and gastric enzymatic degradation which have been lost in both monotreme species^1,2^.

*Nkx3.2* loss-of-function leads to the truncation of the antral portion of the stomach and such mutations also cause a near- or complete loss of antral glandular units in mice^15^. This outcome raises two possible orders of events in which *Nkx3.2* pseudogenisation and gastric gene loss could have occurred in the monotreme ancestor. The first being reflective of the mice mutants in which *Nkx3.2* pseudogenisation causes a loss of the majority of G cells, a disruption of the gastrin physiological pathway and subsequent loss of acidic luminal environment and gastric genes. A lack of expression in *Nkx3.2* suggests that beyond the protein-coding component there is no regulatory functionality of the gene and therefore over time there has been an accumulation of additional mutations. This contrasts the complete loss of gastric genes from the monotreme genome which may suggest that *Nkx3.2* became non-functional after the loss of an acidic environment. Therefore, it is more likely that enzymatic degradation became redundant in the monotreme ancestor and genes conferring an acidic luminal environment and gastric enzymatic degradation became obsolete. This would be followed by a redundancy in the function of G cells and subsequent loss of antral glandular units through *Nkx3.2* pseudogenisation.

### Eco-evolutionary adaptation of the monotreme ancestor stomach

The pseudogenisation and loss of gastric genes and morphology in the monotreme ancestor is also present sporadically in aquatic gnathostome lineages. Select teleost species (e.g. *Takifugu rubribes*, *Danio rerio* and *Oryzias latipes*) and Chimaeriforme species (*Callorhinchus milii*) have converged upon an agastric phenotype associated with pseudogenisation or loss of the hydrochloric acid-associated and gastric enzyme genes as observed in the monotremes^5,46^. Lineages have differed in proposed selective pressures for these losses such as rudimentary gas exchange organs (*Hypostomus plecostomus*), swallowing sea water (*T. rubribes*), adaptations to diets high in calcium carbonate (*Anarhichas lupus*) or swallowing large amounts of undigestible material such as in detritivores (*Mugilidae spp.*)^47,48^. Though selective pressures differ, with the exception of the terrestrial short-beaked echidna, the agastric phenotype has only been observed in aquatic/semi-aquatic vertebrate lineages.

Limited fossil, physiological and molecular evidence has led to debate about whether the platypus lineage is associated with a change from a terrestrial to a semi-aquatic life ecological specialisation in the monotreme ancestor or vice versa^49,50,51^. The semi-aquatic platypus forages at the bottom of freshwater ecosystems to store bottom-dwelling aquatic invertebrates and gravel in their cheeks which have accessory maxillary pads for mechanical digestion of food at the water’s surface^2^. The combination of feeding behaviours is suggestive that potential accumulation of detritus in the stomach may drive loss of an acidic gastric environment such as found in *Mugilidae* species. If the agastric phenotype were a trait associated with aquatic/semi-aquatic organisms then this molecular evidence would support a semi-aquatic monotreme ancestor with subsequent adaptations to the echidna lineage accounting for return and specialisation in terrestrial ecologies.

## Conclusion

Here, we investigated the genetic underpinnings of the monotreme antropyloric region and found that all constituents of the vertebrate antropyloric pathway, with the exception of *Nkx3.2*, were largely conserved and expressed. Interestingly, the functional loss of *Nkx3.2* is consistent with losses of antropyloric segments such as the antral glandular epithelium in both species and the pylorus in the platypus. The discrepancy between the loss of pylorus in platypus and pyloric-like restriction in the echidna may be associated with independent evolution in *Grem1* and *Bmp4* sequences, but regardless leaves the question of whether retention of reinvention of pyloric-like restriction occurred in the short-beaked echidna lineage. We propose that the loss of *Nkx3.2* in the monotreme ancestor led to a subsequent aglandular gastric phenotype in both lineages, and moreover, to the loss of hydrochloric acid production and gastric enzyme genes. The polyphyletic presence of the agastric phenotype in aquatic and semi-aquatic teleost and monotreme lineages, with the exception of the short-beaked echidna, led us to speculate that convergent eco-evolutionary dynamics may be responsible and may offer more evidence for a semi-aquatic monotreme ancestor.

## Acknowledgements

The authors would like to thank Dr Melinda Kulas Jasper (The University of Adelaide) and Emeritus professor Jeremy Timmis for their expertise in PCR amplification and manuscript editing respectively. The authors also thank the late Rory Adelson (The University of Adelaide) for their moral support during the publishing process. Jackson Dann is funded by the Research Training Program.

## Notes

### Competing Interest Statement

The authors have declared no competing interest.

