## Supplementary Table 1 and Supplementary Figure 1 for "Pseudogenisation of NK3 Homeobox 2 (*Nkx3.2*) in Monotremes Provides Insight into Unique Gastric Anatomy and Physiology"

Table S1: Bioinformatic analysis of genes involved in antropyloric development and description of evolutionary changes to sequence structure or canonical motifs likely to affect protein structure, function or processing. Green highlighting of the gene name indicates significant expression (> 0.5 TPM) from any RNA-seq tissues, whereas yellow in the *Pdx1* box indicates ambiguous expression (> 0.5 TPM for echidna, 0 TPM for platypus).

| Gene | Epithelial/ Mesenchymal | Role in Gastric Development | Monotreme Sequence Identities | Gene phylogeny topology and significant sequence changes description |
| --- | --- | --- | --- | --- |
| *Nr2f2* | Epithelial | Epithelial-mesenchymal transition, downstream target of hedgehog and knockout is phenocopy of hedgehog knockout^12^. | Ranged ~ 88-100%. Expected compared to other vertebrate species. | Gene topology reflects known species topology. No significant sequence changes noted in conserved domains^27^. |
| *Pdx1* | Epithelial | Differentiation of G cells, enterochromaffin cells and D cells in antropyloric glands^52^. | Ranged ~ 61-76%. Expected compared to other vertebrate species. | Gene topology reflects known species topology. No significant changes noted in DNA-binding domain. The echidna sequence contains only position 2 P of conserved (D/EPEQD) residues in the PCIF-1 domain. Substitutions include: 1 (D/E > A), 3 (E/Q > P), 4 (Q > P), 5 (D > G)^29^. |
| *Shh* | Epithelial | Smooth muscle differentiation, gut rotation, differentiation of mesenchyme into stomach and pancreas, stomach-intestinal boundaries^10^. | Ranged ~ 75-86%. Expected compared to other vertebrate species. | Gene topology reflects known species topology. Hedgehog signaling domain motifs are conserved. There are large insertions of ~45 amino acids in the hint domain and deletions of ~30 amino acids at the C terminal domain^30^. |
| *Ihh* | Epithelial | Stem cell differentiation and proliferation, differentiation of mesenchyme into stomach and pancreas^10^. | Ranged ~ 52-90%. Expected compared to other vertebrate species. | Gene topology reflects known species topology. There is a large ~83 amino acid insertion in the N-terminal domain for the echidna, part of signaling domain of the peptide^30^. |
| *Sox9* | Mesenchymal | Expressed in pyloric mesenchyme and regulates *Grem1* expression. Mis-expression changed gizzard epithelium to pylorus-like epithelium with ectopic *Pdx1* expression^9,53^. | Ranged ~ 77-98%. Expected compared to other vertebrate species. | Gene topology reflects known species topology. No significant changes noted in conserved domains^31^. |
| *Six2* | Mesenchymal | Regulator of *Nkx2.5*, *Bmp4*, *Sox9* and *Grem1* in pyloric determination. Null mice lack pylorus and stomach mucosa are hypertrophic^40^. | Ranged ~ 92-100%. Expected compared to other vertebrate species. | Gene topology reflects known species topology. No significant changes noted in conserved domains^32^. |
| *Wnt5a* | Mesenchymal | Is downstream from *Nkx3.2*/*Barx1* and inhibition of either result in relaxation of pyloric restriction and ectopic *Wnt5a* signaling in pyloric region^5^. | Ranged ~ 83-100%. Expected compared to other vertebrate species. | Gene topology reflects known species topology. No significant changes noted in conserved domains^33^. |

| *Barx1* | Mesenchymal | Gastric epithelial differentiation from mesenchyme, acts upstream of *Nkx3.2*^9,55^. | Ranged ~ 89-100%. Expected compared to other vertebrate species. | Gene topology reflects known species topology. No significant changes noted in conserved domains^34^. |
| --- | --- | --- | --- | --- |
| *Bmp4* | Mesenchymal | Expression begins in mesenchymal stomach and intestines but terminates at site of pylorus, most likely down-regulated by *Six2* and *Nkx3.2*^,37^. | Ranged ~ 74-90%. Expected compared to other vertebrate species. | Gene topology places monotremes within reptilian taxa. Echidna has missense mutation (A/S > P) in furin S2 cleavage motif (RS/AKR). Both species contain a small insertion (2-5 AAs) before S1 cleavage site (RISR)^37^. |
| *Grem1* | Mesenchymal | Expressed strongly in pylorus mesenchyme, a known BMP inhibitor^56^. | Ranged ~ 63-95%. Expected compared to other vertebrate species. | Sequence identities averaged ~63-95%, in range with other vertebrate species. Gene topology placed monotreme sequences diverging before reptilian species. Monotreme species shared missense mutations in conserved BMP4 inhibition motif (PKKFTTMMVTLNCPELQPPTKKKRVTRXVKQ). Mutations: 2 (K > R), 3 (K > A), 5 (T > S), 12 (N > S), 15 (E > D), 23 (V > F)^36^. Echidna sequence contains 16AA N-terminal insertion. |
| *Nkx2.5* | Mesenchymal | Mesoderm expression distal to stomach correlates to future site of pylorus^5,7^. | Ranged ~ 60-65%, lower compared to other vertebrate species (~75-95%). | Gene topology placed divergence of marsupials earlier than monotremes. Several in/dels and missense mutations outside of conserved domains contributing to low sequence identity. Insertion of 4-6 (GP) repetitive residues in NK2-SD domain in echidna and platypus respectively^35^. |

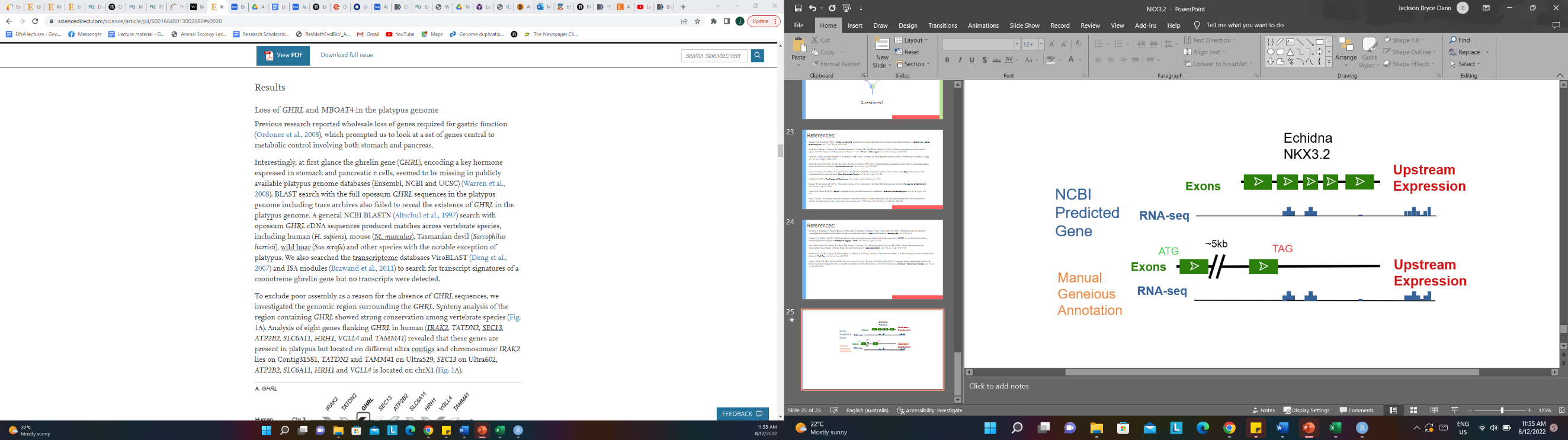

Figure S1: A comparison of the echidna (*T. aculeatus*) *Nkx3.2* gene as predicted by the gnomon algorithm in NCBI GenBank versus the genomic sequence as manually annotated in the Geneious program through comparison to conserved vertebrate sequences and gene structures. Green blocks denote exons, lines are introns and blue blocks beneath sequence structure are expression levels from liver RNA-Seq.
